# Needing: an active inference process for physiological motivation

**DOI:** 10.1101/2022.10.24.513547

**Authors:** J. Bosulu, G. Pezzulo, S. Hétu

## Abstract

Need states are internal states that arise from deprivation of crucial biological stimuli. They direct motivation, independently of external learning. Despite their separate origin, they interact with reward processing systems that respond to external stimuli. This paper aims to illuminate the functioning of the needing system through the lens of active inference, a framework for understanding brain and cognition.

We propose that need states exert a pervasive influence on the organism, which in active inference terms translates to a “pervasive surprise” - a measure of the distance from the organism’s preferred state. Crucially, we define needing as an active inference process that seeks to reduce this pervasive surprise.

Through a series of simulations, we demonstrate that our proposal successfully captures key aspects of the phenomenology and neurobiology of needing. We show that as need states increase, the tendency to occupy preferred states strengthens, independently of external reward prediction. Furthermore, need states increase the precision of states (stimuli and actions) leading to preferred states, suggesting their ability to amplify the value of reward cues and rewards themselves.

Collectively, our model and simulations provide valuable insights into the directional and underlying influence of need states, revealing how this influence amplifies the wanting or liking associated with relevant stimuli.

## 1 Introduction

Our bodies preserve their stability against the many disturbing forces through homeostasis and its mechanisms (Cannon, 1939). Among the mechanisms of homeostasis and its adaptive form, allostasis (Sterling, 1988), there is a particular one that we will refer to as “needing”. Generally, needs intensify, exerting a pervasive influence on various aspects and dimensions of our lives.

For example, when experiencing hunger, its effect persists until satisfied by food. This principle extends to other needs such as thirst and sleep. Therefore, states of need become pervasive, shaping our perceptions, decisions, and more.

Needing is a process related to internal states (Craig, 2003; Livneh et al., 2020) characterized by a deprivation of essential elements crucial for life or survival (Bouton, 2016; MacGregor 1960; Baumeister & Leary, 1995). Here we refer to those (internal) states as need states. Such states (through the needing mechanism) have a directional effect on motivation (Dickinson & Balleine 1994; Balleine, 1992; Bosulu et al., 2022). Needing also interacts with other subsystems that process external stimuli, such as wanting, liking or interception. Thus, needing is both separate from (e.g. see Hogarth & Chase, 2011; Watson et al., 2014), but also interacts with (see Berridge, 2004), different reward related subsystems. This happens in an independent way; for instance, need states tend to amplify the incentive salience of relevant pavlovian cues that generate “wanting” (Toates, 1994; Berridge, 2004) independently of “liking” and (re)learning (Berridge, 2012; 2023). Yet needing can, independently of wanting (i.e. of pavlovian cues) amplify “liking” or pleasure (Cabanac, 2017; Berridge & Kringelbach, 2015; Becker et al., 2019) and influence learning (Dickinson & Balleine 1994; Balleine, 1992; Wassum et al., 2011; Salamone et al., 2018) and interoceptive prediction (Livneh et al., 2020; Bosulu et al., 2022), and needing can also directly activate relevant actions (Passingham & Wise, 2012) or explorative behavior (Panksepp, 2004) as well as behavior related to autonomic and neuroendocrine levels (Swanson, 2000).

Being related to internal states (Sterling & Laughlin, 2015), needing generates the value attributed to rewards “from within”, and is thus separate from the external prediction of reward. Indeed, need states can produce unlearned fluctuations or even reversals in the ability of a previously learned reward cue to trigger motivation/wanting (Berridge 2012; 2023). In the same sense, needing can also modify the perception (e.g. the pleasure) of relevant stimuli independently of their original sensory valence (Cabanac, 2017; Cabanac, 1971). Water tends to acquire a pleasing taste when one is thirsty, and a warm environment or object feels comforting when experiencing cold. Conversely, when too hot, seeking out a cool place can be equally satisfying. These instances highlight how the directional impact of needing plays a crucial role in shaping motivation, ultimately influencing what is perceived as rewarding within a given need state. This directional influence of needing has the ability to alter the perception of pertinent stimuli, irrespective of the predicted or “actual” value of the reward.

Nevertheless, a comprehensive formal framework that encompasses the aforementioned findings and specifically elucidates the following aspects remains elusive: (1) the mechanism by which needing steers motivation, independently of the external world, and (2) the intricate interplay between needing and external rewards processed within other subsystems, such as wanting associated with relevant pavlovian cues; or liking linked to relevant hedonic sensation. In the following sections, we will first tackle these points/questions conceptually, using formal methodologies from active inference (Parr et al., 2022) and introduce the concept of pervasiveness of a need state. Subsequently, we will present two simulations that delve into the functioning of the needing system and its interactions with wanting (and liking).

## 2 Needing system: a conceptual perspective

In the next two sections, we present a conceptual perspective aiming to elucidate two fundamental aspects of needing: (1) its directional impact, and (2) its interplay with other subsystems.

### 2.1 The directional effect of needing

#### 2.1.a In order to remain within their physiological boundaries, organisms are endowed with a-priori preferred states that they tend to occupy

A fundamental objective of organisms is to regulate and maintain their internal states within specific narrow limits (Cannon, 1939; Barrett, 2017; Sterling & Laughlin, 2015; Friston, 2006). For instance, the average normal body temperature for humans is generally between 36.1°C (97°F) to 37.2°C (99°F), which is a very small range compared to the range of possible temperatures in the universe, from the absolute zero to trillions of degrees. The same is true for levels of glucose or the balance between water and salt in the body. The underlying principle is that the spectrum of “states” conducive to life is exceedingly limited in contrast to the immensely large number of alternative combinations that wouldn’t sustain life. Therefore, to ensure a living organism remains within its normal physiological boundaries, natural evolution might have established these boundaries as innate preferred states—possibly encoded genetically—that the organism consistently endeavors to attain. From the formal standpoint of active inference, these preferred states (which might correspond, for instance, to physiological bounds) are referred to as (empirical) priors. They carry a higher probability of realization from the organism’s perspective, meaning they are less surprising (Friston, 2010).

#### 2.1.b Not being within prior preferred states is “surprising” and not having a path (i.e. a relevant reward) to get back to the preferred state creates entropy

Here, the surprise associated with a state, denoted h(y), is the “negative” of being probable and simply means less probable. Also, this surprise is not cognitive as the word “surprise” is commonly used. Anecdotally, for a fish, being out of water would count as a surprising state, as would a very thirsty human. A key claim of active inference is that any self-organizing system must minimize such surprise in order to resist a natural tendency to disorder (Friston, et al., 2006; Friston, 2010) and in the case of our fish, death. Formally, the notion of surprise is closely related to the notion of entropy. Entropy, denoted as H, is the long-term average of the surprise and, here, it expresses the uncertainty related to which state must be occupied. If an organism is endowed with priors about the (preferred) states to occupy, these states have a high prior probability to be occupied from the perspective of the organisms and achieving them reduces the organism’s surprise and its long-term average: entropy (Parr et al., 2022; Friston, 2010). Hence, there is a negative correlation between the availability of a path to the preferred state (i.e. rewards, with their related cues and actions, that lead to the preferred states) and the need induced entropy, because of the prior tendency to visit (more often) those states which results in lower entropy. Importantly, the notion of being in a surprising state (or in other words, being far from preferred states) links well to the concept of “needing” discussed in the Introduction. In the same way being in a surprising state entails an (informational) cost, a state of need entails a (biological) cost if a creature does not respond to the need (see MacGregor 1960; Baumeister & Leary, 1995). When a living organism moves away from its preferred state, it is in a state of “need” - which amounts to having a tendency to occupy (again) the preferred states. Formally, *p*(*Y*|*C*) (Eq. 1)if represents the probability that a state (*y*) should occur, given prior preferences (denoted as *C*), then a state a need state can be represented as:

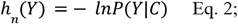

where *h*_*n*_ represents the “need-related” surprise of a sensation or state *y*, which is equal to the negative log probability of being in (or observing) a state *Y* given the distribution of prior preferences, *C*. Note that for simplicity, in this article we will collapse the notions of “state” and of “observation that can be obtained in the state”, which are typically distinct in active inference and more broadly, in Partially Observable Markov Decision Processes.

#### 2.1.c Pervasiveness of need states: need states are pervasive over time and to other states, except to the one (or few) state that alleviates that need

Pervasiveness is the key hypothesis that relates ‘needing’ to prior preferences over states to occupy as well as to the surprise and entropy of not being in such states. By “pervasiveness,” we refer to the potential increase, “as time goes’’, in a need state’s impact on “other states”, i.e. other dimensions of life, unless one transitions to the (only) state that satisfies that need. Formally, we represent the pervasiveness of the need state, or the fact that the need state propagates a negative valence to any other state i of the organism, in the following way:

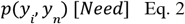

where *y*_*i*_ is any state, *y*_*n*_ is the need state and *p*(*y*_*i*_, *y*_*n*_ ) is the joint occurrence (conjoint probability) of the ith state and the need state. Looking at Eq. 1 it becomes clear that the surprise, noted as − *ln*[*p*(*y*_*i*_ |*C*_*p*_)], gets greater for *almost* all states. So, when the organism is in a need state, all states eventually become surprising, i.e. paired with negative valence, because they can jointly occur with the need state. The only exception is the rewarding state (and the fewer states that lead to it) that alleviates that specific need, because the probability that it jointly occurs with the need state is zero (or close to zero, *eventually*). This allows animals to follow a gradient of surprise minimization. Practically speaking, if one is in a need state, e.g., a state of “hunger”, this need state would propagate a negative valence and (need related) surprise to all the other states (e.g., the states of “sleep”, “run”, “play”, etc.). The only exception to this is the state “having food in the stomach”. Hence, pervasiveness increases entropy (the long-term average surprise) because almost all states become surprising when one is in a need state. The concept of pervasiveness is illustrated in the figure below:

#### 2.1.d Needing induces a tendency to transition, from states to states, towards the preferred state that alleviates it

The perception of a need state translates into a “goal” of reducing surprise by reaching the preferred states, e.g., states that represent adaptive physiological conditions (Friston, 2010). Such a tendency could activate an action or a policy (i.e., a sequence of actions) that compels creatures to seek out the (valuable) preferred states. Note that the actions or policies that resolve a state of need could in some cases correspond to (fixed) regulatory actions, such as autonomic reflexes, as opposed to action courses determined by the circumstances of the external environment (Sajid et al., 2021). With time, the states that the creature occupies when pursuing a policy that resolves a need can become valued per se (Friston & Ao, 2012). In other words, when the creature pursues a course of actions towards the preferred state, all the intermediate states (here intended in a broad sense that encompasses situations, actions, stimuli, etc.) can become valued and needed, through a pavlovian mechanism (Bouton, 2016; Berridge, 2018). For instance, when moving from a state of hunger to a state of satiety, some intermediary states, such as the gustatory stimulus associated to having food and the act of eating, would become valued, because they are in the path towards the preferred (satiety) state (Pezzulo et al. 2015). In sum, a creature that is far from preferred states would experience a need (e.g., for food) – and then start to prefer the valued states (here, intended in a broad sense that encompasses stimuli, actions, etc.) that secure the relevant reward (e.g. food) or are part of the experience of having food (e.g. food cues or relevant actions). In this sense, the tendency to occupy preferred states confers to need states the possibility to influence – and give value to – stimuli or actions that are either costly states (noted *S*(*h*_*n*_)) that lead to surprise, or in the path towards the preferred state (noted π(*p*)). In other words, when one experiences needing, any state (stimulus or action) that is in the path to the preferred state will become something one needs (and hence valued) because it reduces need-related entropy. Hence, the directional effect of need states on motivation could come from the tendency to occupy these preferred states.

#### 2.1.e Needing is an (active) inference process that aims at reducing pervasive surprise

A need state could be defined as a state that is pervasive over time and over other dimensions of the individual’s life, and whose negative impact is surprising with regard to prior preferences. Needing can then be defined as an active inference process that aims at reducing such *pervasive surprise* by inducing a tendency to transition, from states to states, towards the preferred state. The effect of needing does not have to be learned anew, but it is “actively inferred” - reflecting the fact that need states (hunger, thirst) and needed stimuli (food, water) are related to fundamental priors that are potentially sculpted by natural selection. Indeed, even very simple creatures that have limited (or no) learning capacities are able to navigate their environment adaptively to fulfill their current needs, e.g., by following food gradients (Sterling & Laughlin, 2015). The implications of this definition of needing is that it allows shifts in need states to direct motivation towards relevant stimuli even without learning, for instance through alliesthesia (the natural change in sensation/perception of a relevant stimulus induced by a need state). A striking illustration of this phenomenon occurs when animals would suddenly have an increase in “wanting” associated with a salt cue, when depleted of salt even in absence of (re)learning (Berridge, 2012; 2023). This happens even when that cue was learned to be predictive of negative outcome and without the animals having (re)learned about the cue (or the salt) through tasting (and liking) it in the newly induced need (salt depleted) state. Our paper provides a plausible explanation through the lenses of active inference and pervasiveness. The switching from learned negative outcome to wanting happens through pervasiveness which progate surprise to all other states (stimuli, cues, goals, etc) except the states on the path to the priori preferred state. Notably, this category encompasses the (memories of) salt and its cue, as these stimuli still co-occurred with salt in the body, despite being learned to be negative (see Smith & Read, 2022). Thus by reducing such pervasive surprise through an active inference process, the animal would tend to have an increase in wanting associated with such salt cues without (re)learning. This adjustment is achieved through precision increase as elaborated below.

### 2.2 Interaction between needing and other subsystems

#### 2.2.a Needing modifies perception of rewarding states processed within subsystems through increase in precision: such precision is signaled by neurotransmitters of each subsystems

The effect of needing on wanting (and on other phenomena such as pleasure and liking) could be conceptualized by appealing to the formal notion of *precision* in active inference. Mathematically, precision is a term used to express the inverse of the variance of a distribution which in our context can be seen (loosely speaking) as the inverse of entropy (Friston, 2010; Holmes 2022) – in the sense that the higher the entropy, the lower the precision. In predictive coding and active inference, precision acts as a multiplicative weight on prediction errors: prediction errors that are considered more precise have a greater impact on neural computations (Parr et al., 2022).

With respect to need states, precision can be interpreted as a higher salience attributed to the most relevant stimulus given the need state. There are different precisions associated with different subsystems, such as those related to interoceptive streams, rewards, policies, etc. (Parr et al., 2022). At the neurophysiological level, policy precision, or the confidence that a policy should be followed, is typically associated with the dopaminergic subsystem in active inference (FitzGerald et al., 2015; Parr et al. 2022, Holmes, 2022). Therefore, pavlovian cues that enhance policy precision and confidence that the policy should be followed would trigger dopamine bursts, which will attribute incentive salience, i.e. wanting, to such cues (Berridge, 1996; Berridge, 2007). This is in line with the idea that dopamine is linked with incentive salience and wanting; but also with reward cues and behavioral activation as they typically co-occur (Hamid et al., 2016, but see Berridge, 2023). Rather, precisions regarding hedonic contact with the reward (to ask questions such as: is it good?) or the state of satiety (to ask questions such as: am I well?) might be mediated by the opioid subsystem (Berridge & Kringelbach, 2015) and the serotonin subsystem (Liu et al., 2020; Luo et al., 2016; Parr et al., 2022). The impact of needing on stimuli that are processed within these subsystems are “natural” increases in precision due to prior preferences. The need-induced increase in precision implies more certainty that the state to which the stimulus or policy leads to is the least surprising. It is this certainty that amplifies wanting, liking, etc. and it is an active inference process that may be separated from learned reward prediction (also see Berridge, 2023).

This discussion helps appreciate the deep interdependence between needing (which acts on the system as a whole) and wanting as well as other subsystems such as the hedonic/liking and interoceptive ones. When one is in a surprising (need) state, the presence of a cue (e.g., a traffic or restaurant sign) might reduce uncertainty about goal/reward achievement by improving policy precision via the dopamine subsystem (wanting). The presence of, or contact with, the reward itself might reduce entropy by enhancing precision through the opioid subsystem (pleasure/liking). Finally, being in a preferred state or moving towards the preferred state determines an increase of precision - which is due to the fulfillment of prior preferences - via the serotonin subsystem (well-being).

In all these cases, rewards and cues are useful information sources: they reduce entropy by signaling the availability of a path to preferred states (π(*p*)), or equivalently a path away from surprising states (*S*(*h*_*n*_ )), given some prior preference (*C*_*p*_ ). Indeed, from the point of view of the organism in a need state, both encountering either a relevant reward or a cue that leads to that reward would reduce uncertainty about how to move to the preferred state and alleviate the need. This is despite making contact with a reward and encountering a cue that predicts reward might be treated in different subsystems: the liking and the wanting subsystems, respectively.

### 2.6. Summary

Our discussion so far has highlighted two crucial aspects of needing and its interactions. First, need states exert a directional influence on choices separately from reward prediction. This is due to the pervasiveness of need states that make states surprising with regard to prior preferences, and to the animals tendency to reduce surprise, which naturally increases the value of cues and actions that lead to the preferred state. This tendency exists irrespective of reward prediction (Berridge, 2023; Zhang et al., 2009; Smith & Read, 2022). Second, when there is a path to the preferred state, such as a reward or a pavlovian cue, needing would increase the value of reward or pavlovian cues within the subsystems that process them. This translates into a lowering of entropy about which state to occupy (i.e. entropy of the probability distribution of the to-be-reached states) in order to transition to the preferred state, and thus an increase in the precision of relevant stimuli (e.g. liking) or goal-achieving policies (e.g. wanting). Such high precision could be viewed as (need induced) salience which, for instance, in the wanting subsystem translates into higher incentive salience. With these insights in mind, we now shift from a conceptual discourse to the practical implementation of needing through an active inference framework.

## 3. Mathematical formulation of needing

*In the following section, we present the mathematical formulation by introducing an agent endowed with needing. This agent is utilized in simulations to illustrate the functionality of the needing system and its influence on reward subsystems, such as wanting and liking. In this section, we adopt a parallel structure to that of Section 2. Concerning the directional impact of needing (2.1.), we delve into mathematical specifics of the subsections (from 2.1.a to 2.1.e). Additionally, with respect to the interaction between needing and other subsystems (2.2.), we explore the mathematical intricacies associated with both the need-related entropy, and need-induced precision increase. We then map the mathematical formulation to the conceptual perspective in 3.3 and 3.4*.

### 3.1. The mathematical details of the directional effect of needing

#### 3.1.a The probability of a state given prior preferences

Animals possess inherent prior preferences that condition the probability of what specific states “should” occur, noted as:

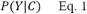

which means the probability of observing a state *Y* (remember that in our setting, hidden states and observations are the same) given prior preferences *C* (i.e. biological costs or rewards).

#### 3.1.b The main formulation of need state as surprise

A need state could be defined as a state that is pervasive over time and over other dimensions of the individual’s life, and whose negative impact is (biologically) surprising with regard to prior preferences. The general notation of such surprise is:

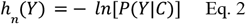

where *ln* denotes a natural logarithm and (*P*(*Y*|*C*) the probability of observing a state *y* given prior preference (C).

#### 3.2.c Formulation for the pervasiveness

The conditional probability of the states *Y* given prior preference will decrease proportionally to the amount of need (*n*) they embed, and this decrease in probability will increase the surprise of being in those states. Due to the pervasive effect of a need state, the value of any state *y*_*i*_ with regard to (the amount of) need (noted *y*_*i,n*_) that one has while being in that state depends on its co-occurrence, i.e. joint probability, with the state that generated the need, noted *y*_*n*_ .

Thus, any state *y* will have a need equivalent to:

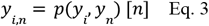

The states that are affected by the need (*n*) will become elements of the set of surprising states, noted *S*(*h*_*n*_ ). Those that are not will be elements of set of states on the path to the preferred state, noted π(*p*). This (i.e. Eq. 2) is to be considered over time as states on the path to the preferred state (such as relevant reward cues) can for a few moments co-occur with that need, but a few moments later, that need state would disappear (as they led to the preferred state). Hence their joint probability with the need state, taking time into account, would still be lower.

We can then note the need-related surprise of each individual state as given the prior preference as :

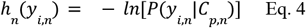

where *C*_*p,n*_ represents the specific priori preferred state (*p*) that alleviates the specific need (*n*).

Assuming that animals have prior preferences over levels of satiety/need (e.g. a preferred level of sugar in the body), the prior preference probability over satiety is equal to p(*C*_*p,n*_) = 1.

Thus, the preference for each state i given the prior preference and the need becomes their joint probability:

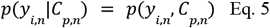

Thus our Eq. 3 can, under those conditions, be written:

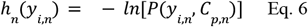

This illustrates that the co-occurrence with a need state is inversely related to co-occurrence with the preferred state. Thus, for a given need, the need-related surprise of each state is the negative logarithm of its joint probability with the preferred state.

Given the pervasiveness of the need state, almost all of the *y*_*i*_ become surprising because their joint probability, i.e. co-occurrence, with the preferred state decreases. Hence, only the states “food in stomach” and the states that lead to it have high probability as they (eventually) co-occur with the priori preferred state. Crucially, as it will become clear in our first simulation, this probability increases as need increases, because the sum of probability of *p*(*y*_*i,n*_|*C*_*p,n*_) has to sum to 1.

#### 3.1.d Formulation of surprise minimization

Since the creature expects to occupy (or to move towards) these a-priori probable states, the prior over states also translates into priors over actions or action sequences (policies) that achieve such states. In this simplified setting, action (and policy) selection simply corresponds to inferring a distribution of states that it prefers to occupy and policies to reach (sequences of) these states. In other words, the active inference agent tends to select policies that lead it to achieve goal states - which in Bayesian terms corresponds to maximizing model evidence (Parr et al 2022).

More formally, under the simplifying assumptions discussed above, the creature strives to maximize a measure of (log) evidence, or alternatively minimize the associated surprise − *ln*[*P*(*y*_*i,n*_ |*C*_*p,n*_)]. The creature does so by minimizing the expected surprise of every state, given the prior preferences, as denoted below:

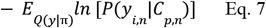

The *E*_*Q*(*y*|π)_ part means that the probability of states/outcomes is averaged across all policies/paths (π).

In active inference, the quantity shown in Eq. 7 *E*_*Q*(*y*|π)_ *ln*[*P*(*y*|*C*)] (without the minus sign) is typically called a “pragmatic value”, and in this setting (with a minus sign) it corresponds to the expected free energy G(π) (an upper bound on expected surprise):

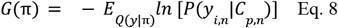

For completeness, it is important to consider that the quantity shown in Eq. 8 (without the minus sign –the “pragmatic value”–is only one of the two terms of the expected free energy G(π) of active inference; however, in our setting there is no ambiguity and the second term (“epistemic value”) is zero.

The expected free energy G(π) is particularly important since it is used for policy selection. Specifically, active inference agents are equipped with a prior over policies, denoted as *P*(π). The greater the expected free energy that policies are expected to minimize in the future, the greater their prior, i.e.

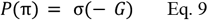

where σ represents the softmax function, bounded between 0 and 1, and enforces normalization (i.e. ensures that the probability over policies sums to one).

#### 3.1.e Mathematical definition of needing from an active inference perspective

Needing is thus the minimization of *G*(π), or to expressing this in a “positive” way, needing is the maximization of *E*_*Q*(*y*|π)_ *ln*[*P*(*y*_*i,n*_ |*C*_*p,n*_)]. In other words, needing induces a tendency to seek states in which the expected log-probability given the prior preference is maximized under that specific (pervasive) need state.

### 3.2. Mathematical detail of the interaction of needing and other subsystems

#### 3.2.a Formulation of precision increase

Now, we consider the effect of a need state on (1) the entropy over the states that a creature plans to occupy in the future by following its inferred policy and (2) the inverse of the entropy, i.e., the precision, which is a measure of certainty about which states to occupy next.

The “need-related entropy” (noted *H*_*n*_ or *H*_*n,p*_, explained further below) is the average, i.e. the expectation (*E*), of all negative log probability of all states Y given prior preference under a specific need, and is given by the formula:

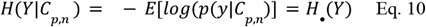

As discussed, given pervasiveness (i.e. joint occurrence with the need state), this variable Y can indeed be a surprising state, i.e. embedded with need states, or it can be a rewarding state, i.e. a state on the path to the preferred state. Thus, the creature’s “need related entropy” (or simply entropy) as follows:

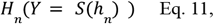

when there is no path to the preferred state, i.e. no reward (in a broad sense); and

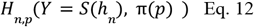

when the cue/reward is present and the creature has a potential path towards the preferred state.

Here, H (i.e. *H*_*n*_ or *H*_*n,p*_ ) denotes the entropy and it can be calculated on two sets of states. When a reward (or its cue) state is available, the entropy is over which states to occupy by the creature when some of the states are on a path π(*p*) to the preferred state (*p*) while the rest of states are not (*S*(*h*_*n*_)). Alternatively, when there is no reward, then the entropy is over which states *S*(*h*_*n*_) to occupy.

Thus, *Y* = *S*(*h*_*n*_ ) means *Y* is any *S*(*h*_*n*_), and *Y* = *S*(*h*_*n*_), π(*p*) simply means *Y* can be any *S*(*h*_*n*_) or any π(*p*). The π(*p*) and *S*(*h*_*n*_) represent states in different subsets of prior preferences, with the π(*p*) representing states that are on the path to the preferred state. These can be viewed as rewarding (or cues) states or events that lead (transition) to the preferred state if one follows a policy leading to the preferred state (discussed in section 3.3). The *S*(*h*_*n*_) represent the states that lead to surprise as they are (eventually) embedded with the need. This formulation highlights that the (need-related) entropy of an agent that faces costly/surprising states *S*(*h*_*n*_) is reduced when there is a path towards the preferred state π(*p*). Thus, the inequality below holds:

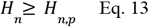

We can calculate the precision as the inverse of the entropy:

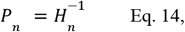

when there is no reward (cue) state, and

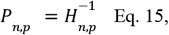

when there is a reward (or cue) state and hence a path to the preferred state.

Given the inequality in Eq. 13, when many states become more surprising in terms of need, i.e. biologically costly, the precision increases, providing that there is a path towards the preferred state, which implies that:

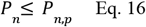

Given that we are discussing the motivational, i.e. active part, here entropy means (average) uncertainty over which state to occupy rather than uncertainty over what state is. Similarly, precision means *certainty over what state to occupy*. The principle is the same whether applied to what states to occupy or what policy to follow. The idea is to make it general so it can apply to incentive salience (wanting subsystem) or to hedonic sensation (liking subsystem), and also to simpler organisms that might not have a sophisticated brain.

In the specific context of the interaction between needing and “wanting” we can draw a connection between our precision equation and the computational models pertaining to wanting and incentive salience proposed by Zhang et al. (2009) and Smith & Read (2022). In both models, a variable denoted as “k” represents the impact of physiological and other brain states on the dopaminergic state. This “k” factor alters the reward value “r,” subsequently leading to an amplification of the reward cues, expressed as the function 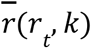. When such a variable “k” is influenced by need states, the function 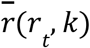 becomes associated with precision over policies that guide the attainment of the preferred state, denoted here as *P*_*n,p*_ .

### Illustrating the connection between needing and wanting

### 3.3 The general and directional motivation of needing

A need state could be defined as a state that is pervasive over time and over other dimensions of the individual’s life, and whose negative impact is (biologically) surprising with regard to prior preferences. Such surprise is noted: *h*_*n*_ (*y*) = − *lnP*(*y*_*i,n*_ |*C*_*p,n*_). The pervasiveness of need states, given into prior preferences, propagates such surprise depending on the joint probability between a need state *y*_*n*_ and another state *y*_*i*_ given by *p*(*y*_*i*_, *y*_*n*_ ) [*n*]. Prior preferences then “judges” how any state *y* should be preferred given the preferred state in which that alleviates need, noted *C*, and in which the system expects to be in, and this is given by: *P*(*y*_*i,n*_ |*C*_*p,n*_ ). Based on that, the pervasiveness of need states make many states less probable, i.e. surprising, noted *S*(*h*_*n*_). Hence the system, expressing states in terms of surprise, i.e. − *lnP*(*y*_*i,n*_ |*C*_*p,n*_), can “infer” the trajectory, i.e. the couple of states which are noted π(*p*), that lead to the preferred state, by choosing the actions from which one expects minimization of such surprise: − *E*_*Q*(*y*|π)_ *ln* [*P*(*y*_*i,n*_|*C*_*p,n*_)]. Needing is this active inference process that aims at reducing such *pervasive surprise* by seeking states that maximize *E*_*Q*(*y*|π)_ *ln* [*P*(*y*_*i,n*_|*C*_*p,n*_)], i.e. states whose expected log-probability given the prior preference is maximized under that specific (pervasive) need state.

### 3.4 Interaction between needing and other subsystems such as wanting, liking, etc

Need states can amplify wanting or liking and thus assign a high precision *P*_*n,p*_ to pavlovian cues or hedonic contacts with rewards that lead to the preferred state. Indeed, by following policies that minimize − *E*_*Q*(*y*|π)_ *ln* [*P*(*y*_*i,n*_ |*C*_*p,n*_ )] towards the preferred state and away from surprising states *S*(*h*_*n*_), when there is a possibility/path to reach such preferred state, the states on that trajectory π(*p*) become more and more probable given the prior preference. This is because, due to pervasiveness, the states on π(*p*) have a lower joint probability with the need state, and this decreases entropy over which state to occupy. In other words when one is in need (surprising) state that becomes pervasive, and there is a path to the preferred state, the entropy *H*_*n,p*_(*Y* = *S*(*h*_*n*_), π(*p*) ) = *H*_*n,p*_ decreases by following the policies that minimize − *E*_*Q*(*y*|π)_ *ln* [*P*(*y*_*i,n*_|*C*_*p,n*_)]. Thus, the inverse of that entropy 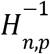, i.e. the precision (or neuronal gain) *P*_*n,p*_ assigned to states (stimuli/events/reward cues) π(*p*) that leads to the preferred state is increased. This increase in precision happens within any subsystem (wanting, liking, interoception, etc.) if such π(*p*) states (stimuli/events/reward cues) happen to be processed by that subsystem. So, the interaction between needing and, for instance, wanting happens when needing enhances mesolimbic dopamine reactivity which assigns higher precision to pavlovian cues that are relevant under the need state. The enhancement of dopamine reactivity amplifies/generates ‘wanting’ associated with relevant pavlovian cues by acting as neuronal gain expressed as *P*_*n,p*_.

## 4. Simulation results

In the next sections, we subsequently illustrate the functioning of the model in two simulations, which exemplify how being in need states influences the tendency to reach preferred states independently of reward prediction (Simulation 1), and how the simultaneous presence of a state of need and the presence of a path to the preferred (reward or goal) state implies low entropy and high precision over which state to occupy (Simulation 2).

### 4.1. Simulation environment

We designed a 3x3 grid-world with 9 states, in which only one state (state 2) is rewarding/preferred, one is a need state (state 8), i.e. it is (biologically) costly/surprising; and the other seven states are “neutral”; see Figure 1 and 2. We can draw a parallel between the state 2 and 8 and human physiological states, such as hunger or temperature: the preferred state (2) corresponds to the optimal interval of sugar level in the bloodstream, or the temperature range (between 36.1°C (97°F) and 37.2°C (99°F)), and the state 8 is a deviation from that.

**Figure 1.**
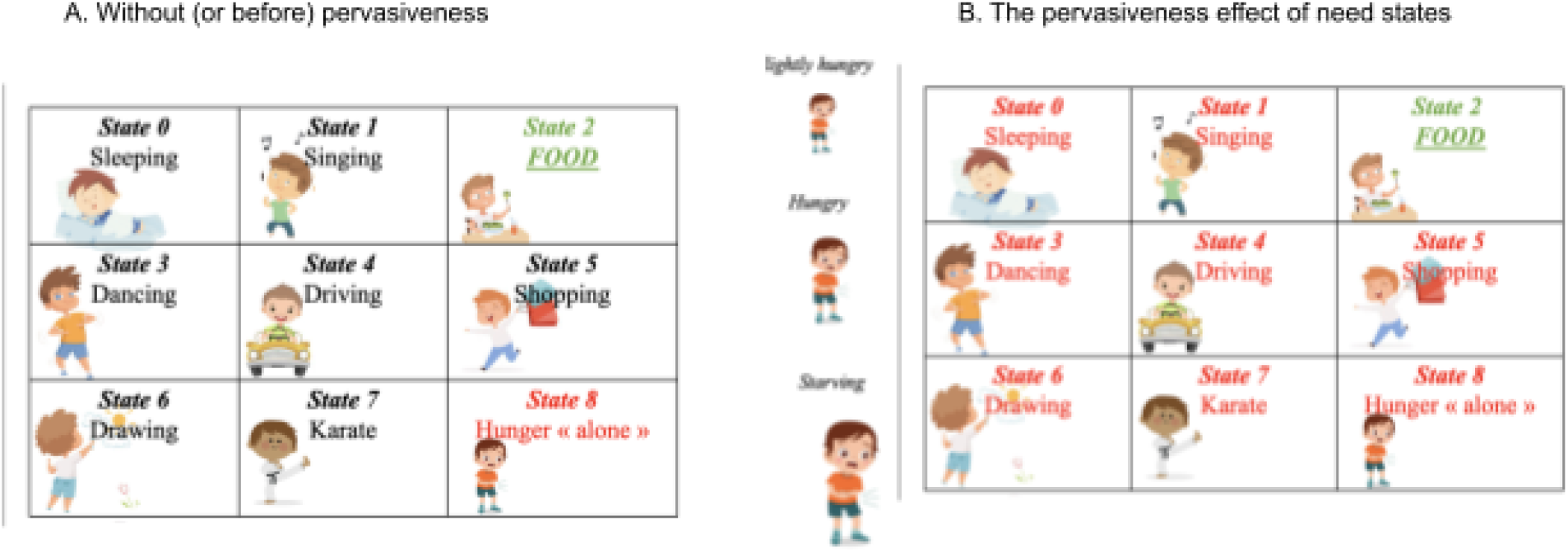
The impact of pervasiveness of a need state on other states. The expansion of the “red” illustrates the pervasiveness effect. In A there is no pervasiveness and hunger is more like an aversive state (state 8). In B pervasiveness occurs and hunger impacts all other states, except the (preferred) state that alleviates it. This increases preference for stimuli/events or actions that lead to the preferred state because there is no other choice. By doing so, it increases the precision (neuronal gain) or weight assigned to relevant stimuli/events or actions within the subsystems that process them.

**Figure 2.**
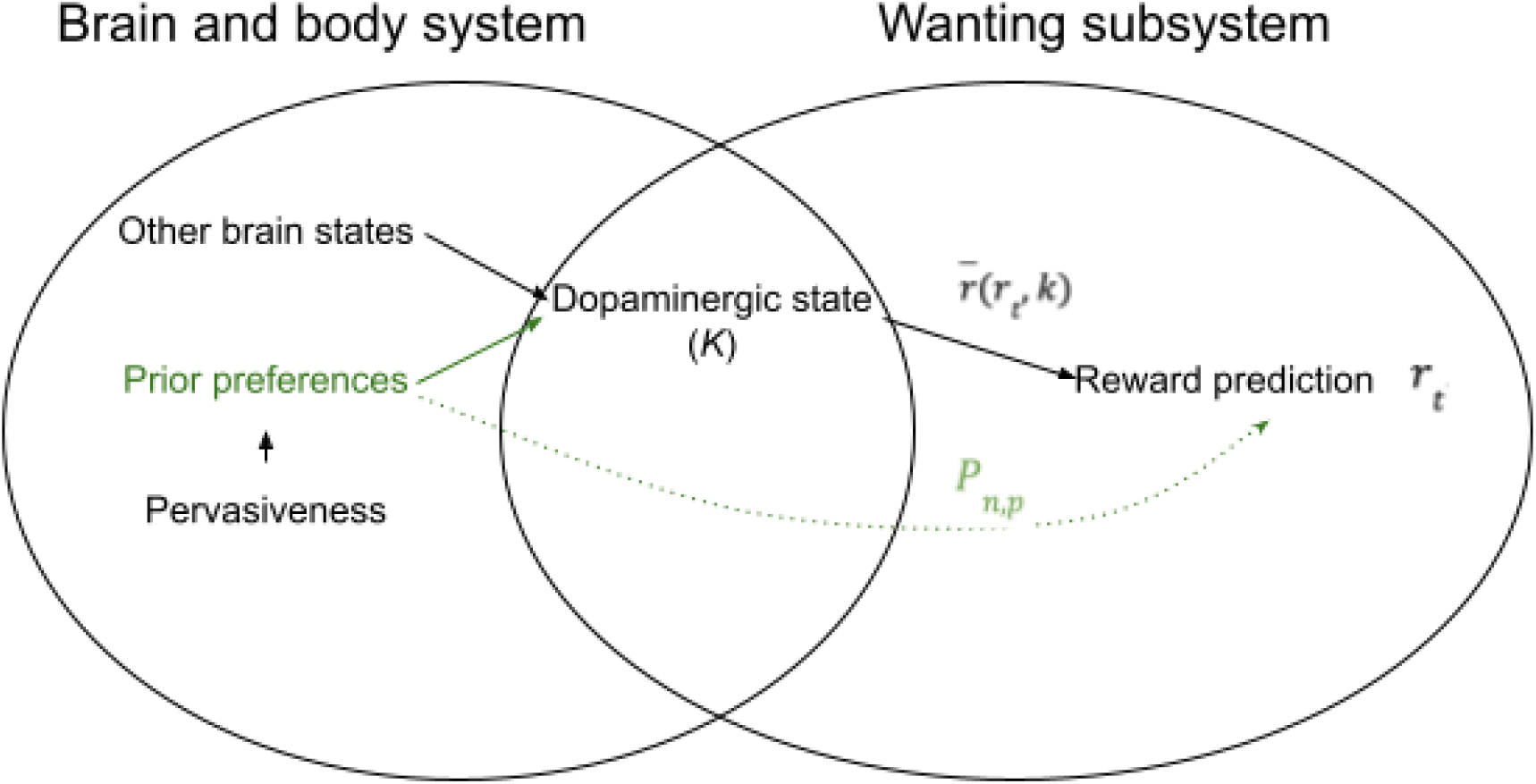
Illustration of the wanting subsystem which depends on the pavlovian cues, paired with some stimulus, noted r_t_, under the influence of different inputs that elevate dopaminergic state k, such that the overall effect is 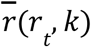 as described by Zhang et al. (2009) and Smith & read (2022). If the input is the need state (in green) then 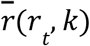 becomes P_n,p_ and needing amplifies wanting.

In the grid-world, each state corresponds to a box in Figure 2. In our simulations below, we will assume that the agents are in a need state (e.g., hunger) that can be low, mild or severe. Accordingly, a negative reward of -1 (slightly hungry), -2 (hungry), or -5 (starving) is assigned to the “costly” state (state 8). State 2 represents a reward/preferred state that gives a reward of 1 and resolves the need state. Note that the agents that dwell in the simulated environment sense this biological reward/cost and will have to compute the *expected* biological costs, or values, by themselves. Of note, the gridworld is used here for visual presentation purposes and the proposed framework can indeed be generalized beyond a gridworld.

### 4.2. Simulation 1: Directional aspect of needing separately from reward prediction

Our simulations will consider two agents (see APPENDIX: SIMULATION AGENTS). Agent 1 incorporates the main ideas discussed in this article about the needing system. It is a simplified active inference model that bases action selection on prior preferences about states and policies (Parr et al., 2022). Agent 2 is used as a comparison, to show that reward prediction alone is not sufficient to explain the directional effect of needing and its interplay with other subsystems such as wanting. It is a simplified reinforcement learning system that based action selection on learned action values computed by reward prediction (Sutton and Barto, 2018) and perceives a “need” without the pervasiveness of need states as described here. The goal of Simulation 1, illustrated below, is to assess the effects of increasing need states on the action selection mechanisms of the two agents.

The results illustrated in Figure 4 show that increasing the cost of the need state, when pervasiveness occurs, significantly increases the probability assigned to policies that reach the rewarding state in Agent 1. This is evident when considering that the probability increases from (about) 0.5, 0.8 and 1 in the three left rows. However, increasing the costs of the need state without need related pervasiveness does not affect reward prediction as shown in Agent 2. This is evident when considering that the softmax of state-action (Q) values assigned by Agent 2 to the rewarding state is always relatively the same in the three right rows. These results help illustrate the idea that costly or need states might exert directional effects and impact on the probability (or tendency) to reach preferred states through pervasiveness, irrespective of reward prediction.

### 4.3. Simulation 2: How needing amplifies wanting

In Simulation 2 (using the same grid-world environment), we ask if being in a state of greater need increases precision associated with rewards, i.e. states that are on the path to the preferred state. We compare two conditions; namely, a condition when a reward is present (i.e. the reward state 2 is baited with a reward of 1) or absent (i.e., the reward state 2 has the same cost as all the other neutral states). The results of the simulations of entropy and precision can be appraised graphically in Fig 5. These results shown indicate that compared to the case with no reward, the condition where a reward is present implies a significant decrease of the entropy over which states the Agent 1 plans to occupy in the path to the reward (Fig. 3, left) and a significant increase of its precision, which is a measure of certainty about which states to occupy in the future (Fig. 3, right). This is because the availability of a reward makes the agent more confident about the states to occupy and the policy to select, whereas in the absence of a reward, all states are equally costly/surprising and the agent has no strong preference about which states to occupy (i.e., high entropy and low precision). The presence of a reward (which in this simulation is known by the agent) is a path that makes it possible to pursue a preferred course of action towards the preferred state, reducing the entropy about the states to occupy and increasing the certainty (precision) about the states to visit in the path towards the preferred state.

**Figure 3.**
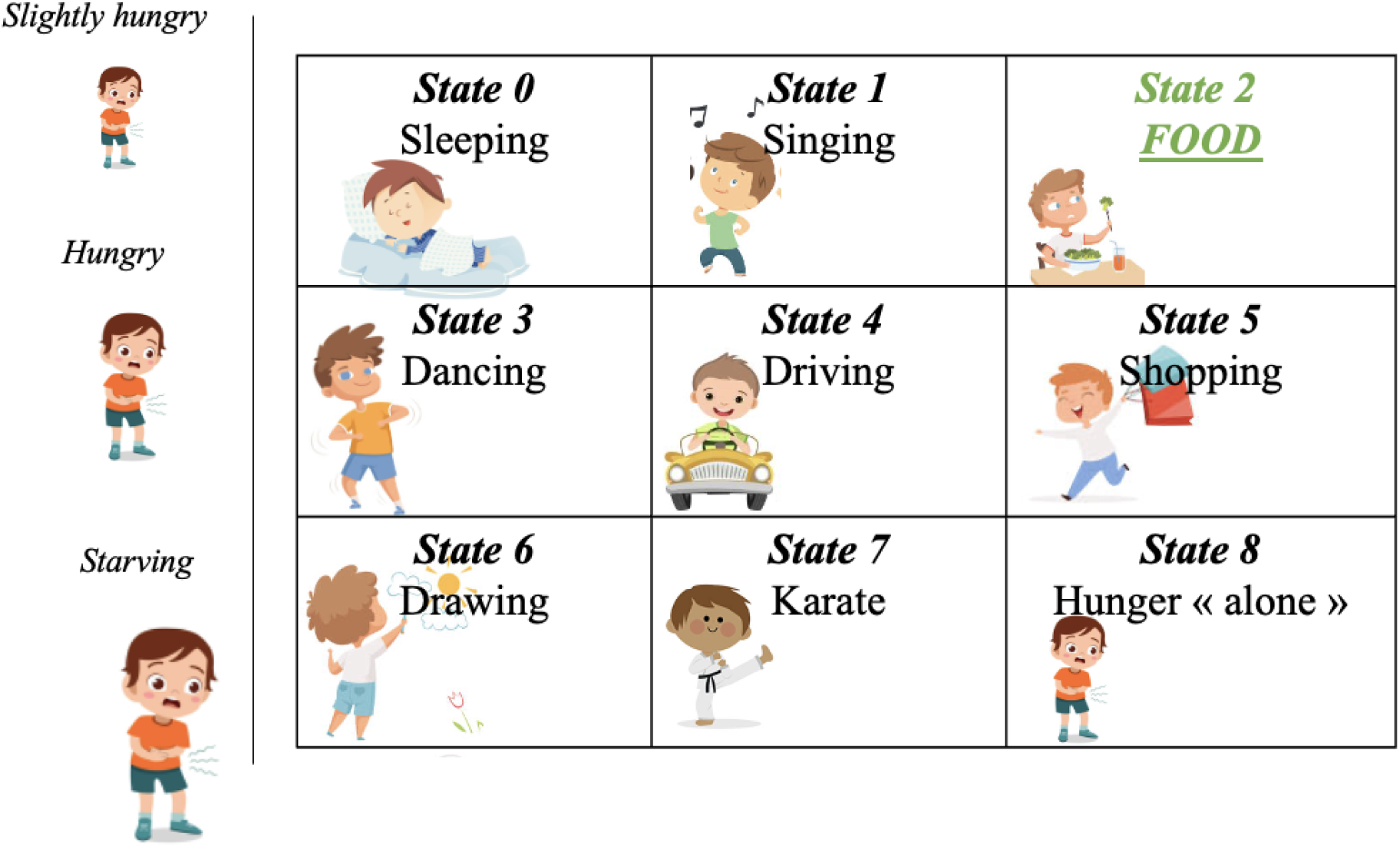
Grid world environment used in our simulations. Each box represents one state in which the agent can be. These include seven neutral states (states 0, 1, 3, 4, 5, 6, 7), a reward state (state 2), and a costly state (state 8). The value of these states is initially unknown. The boxes represent different states and the images on the left represent some increase in hunger. Please note that we put hunger “alone” in order to distinguish between the presence and absence of the pervasive effect. In this case, the pervasive effect of hunger would attain all the other states, except the state 2: food.

**Figure 4.**
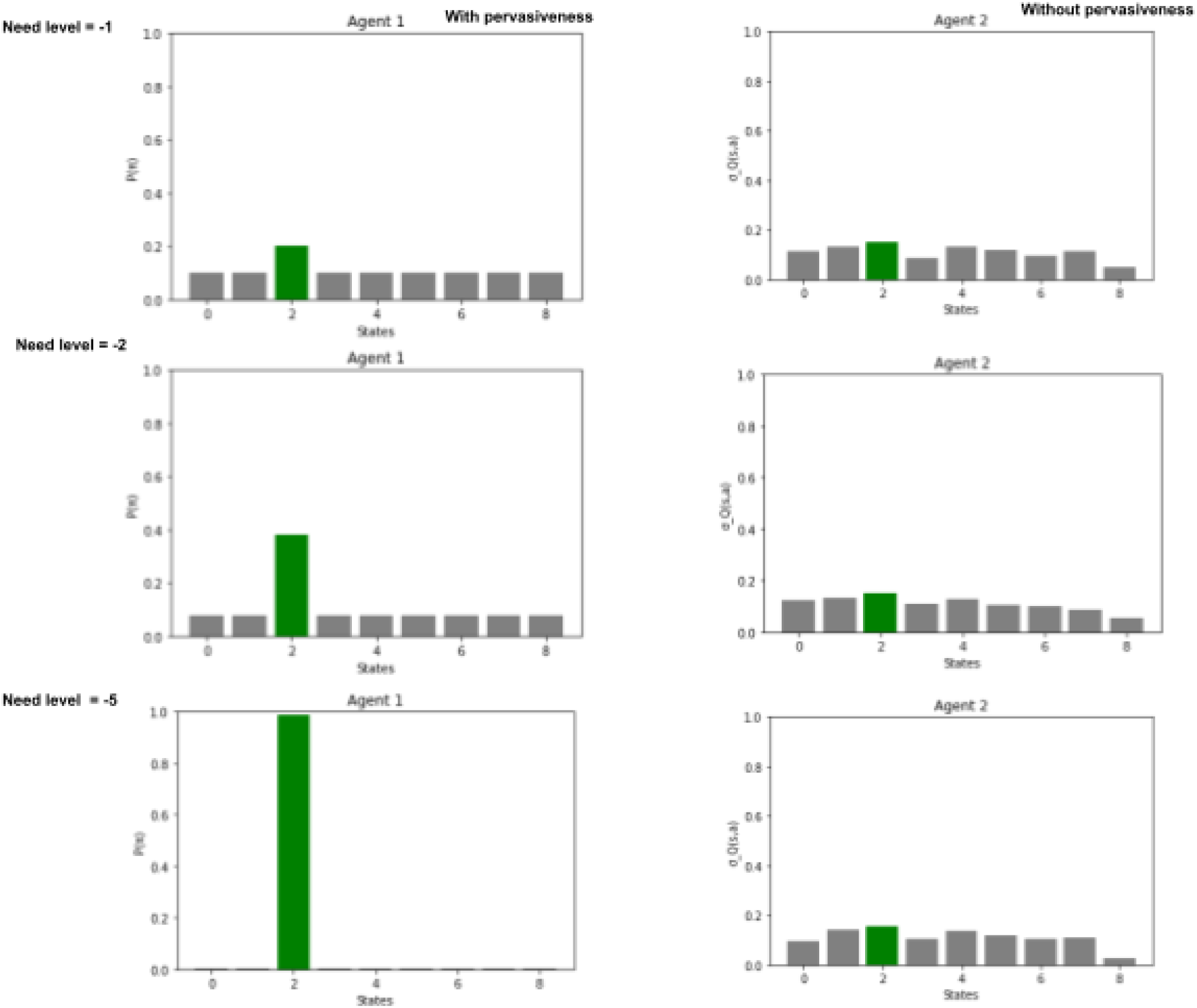
Effects of biological needs on policy selection under pervasiveness and prior preference (Agent 1, left panels) versus computed reward prediction (Agent 2, right panels). The left and right panels show the results for Agent 1 and Agent 2, respectively. For Agent 1, which uses prior preferences, the y axis plots priors over policies P(π) to reach each of the states of the grid-world, whereas for Agent 2, which computes reward predictions, the y axis plots the softmax of the maximal state-action (Q) values (of the action that reaches that state). As evident when looking at the green bars, both Agent 1 and 2 assign the greater probability to the policy (or action) that reaches the rewarding state 2. The three rows show the effects of setting the costly state (state 8, see Fig. 1) to -1, -2 and -5, respectively. The results show that when need states are pervasive, increasing biological needs (across the three rows) increases the probability that Agent 1 selects policies to reach the preferred state 2, but does not increase per se the (external) reward prediction probabilities assigned by Agent 2 to state 2. This can be apprehended by noticing that in the three panels, Agent 1 assigns different probabilities to the policies that reach state 2, whereas Agent 2 assigns (closed to) the same probability to state 2. This is consistent with the idea that need states are pervasive and this allows them to (directionally) influence tendencies (i.e. probabilities) towards the preferred state independently of reward prediction. See the main text for explanation.

**Figure 5.**
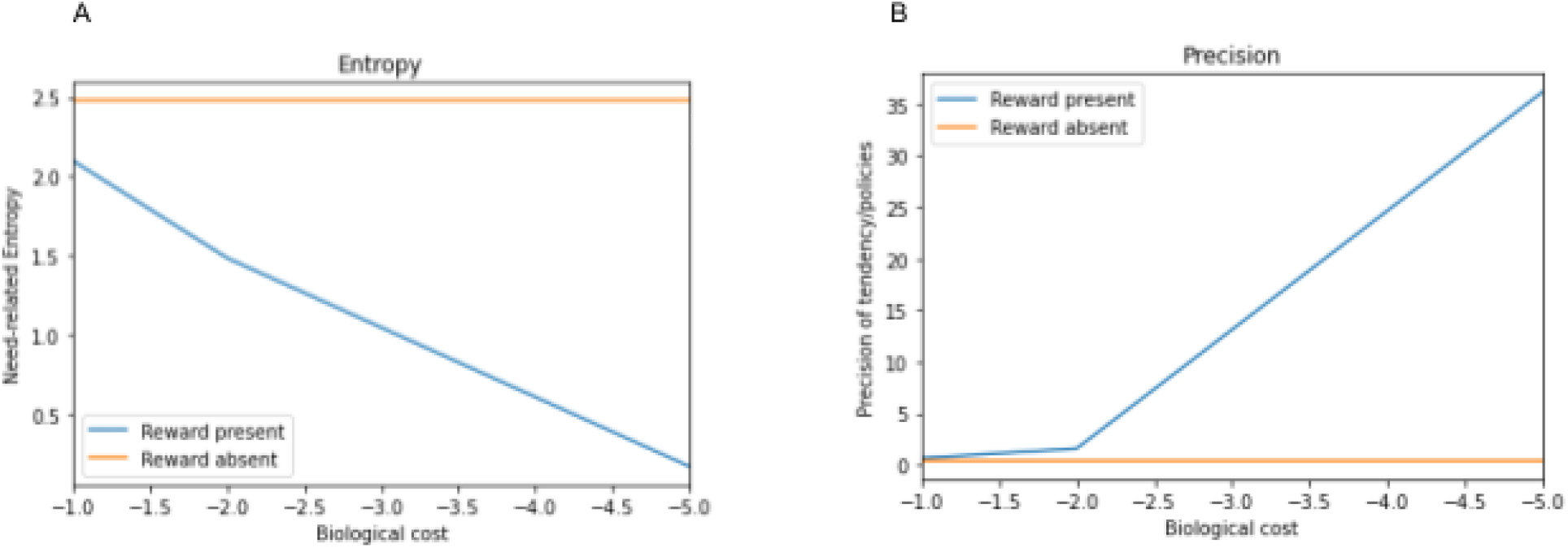
The impact of different need states on the entropy (left plot. A) and on its inverse, the precision (right plot. B), over which state to occupy for an agent that embodies prior preferences, in conditions in which a reward is available (blue lines) or no reward is available (orange lines).

## 5. Discussion

The role of needing and the way it interacts with other reward subsystems are not completely understood. Here, we aimed to provide a computationally-guided analysis of the mechanisms of needing from the normative perspective of active inference and the hypothesis of (embodied) pervasiveness of need states.

We firstly defined a need in terms of pervasiveness and surprise; i.e. a state that is pervasive over time and over other dimensions of the individual’s life, and whose negative impact is surprising with regard to prior preferences. We then defined needing as an active inference process that aims at reducing such *pervasive surprise* by inducing a tendency to transition from states to states, towards the preferred state. Here, the term “surprise” is used in the sense prescribed by theories like predictive coding and active inference. The key idea is that living creatures strive to remain within tight physiological boundaries (i.e. preferred states) and are surprised outside them - like a fish out of water. This perspective suggests that being in a need state is intrinsically costly (where the cost is related to surprise); and thus a state of need may exert a directional effect on action selection and motivation, because creatures would have an automatic tendency to select policies that avoid surprises and lead to preferred states. Importantly, this automatic tendency would be present independently of any reward or reward cue. Moreover, the pervasive effect of a need state acts on the system as a whole, making all states surprising with regard to prior preferences, except the relevant rewarding states, i.e. those on the path to the preferred state. Hence it is the embodiment of that pervasiveness into prior preferences that allows needing to activate prior policies that direct the system, in a gradient-like manner, towards the relevant rewarding states.

We further defined the interaction between needing and other subsystems, specifically the wanting (or liking) subsystem, where needing increases the precision of relevant stimuli or action processed within those subsystems. In the case of wanting, this precision is that of policies that achieve preferred (goal or reward) states in active inference, consistent with previous work that linked policy precision with dopaminergic reactivity (FitzGerald et al 2015, Friston et al., 2017) and thus incentive salience (Berridge, 2004). From this perspective, pavlovian cues that signal that there is a path to preferred state acquire incentive salience (Berridge 2004) and generate “wanting”. Needing amplifies such wanting, as a state of greater need can amplify policy precision by amplifying the value of reward cues that do not co-occur with the pervasive surprise (because they lead to the preferred state). Hence, the higher the initial state of need, the greater the wanting associated with pavlovian cues related to relevant rewards.

### 5.1. Simulation 1: The needing system and its directional effect on behavior

This simulation demonstrates that for an organism that uses prior preferences to embody the pervasiveness effect, need states can have directional effects independently of reward prediction. Needing governs directional motivation due to the inherent tendency of living beings to move towards preferred states. This propensity activates policies leading to those preferred states, which in turn elevates the preference (and value) of states within their trajectory. Homeostasis and allostasis, which assist animals in maintaining viable physiological boundaries (Cannon, 1939; Sterling, 1988, 2004; Barrett, 2017; Holmes, 2022; Demekas et al., 2020), mediate this tendency. Indeed, from the active inference perspective, a living organism continuously strives to reach or remain in its preferred states (which could be sometimes evolutionarily defined, though homeostatic or allostatic regulation, at the somatic, autonomic and neuroendocrine levels (Parr et al., 2022; Swanson, 2000)). These preferred states act as drives or goals that, via homeostasis and allostasis, direct action (Barrett, 2017); hence the directional effect of need states. This directional influence, contingent on the propensity to occupy preferred states, also underlies the amplifying effect of need states on wanting, liking/pleasure, interoceptive prediction, choice, etc., by increasing the precision of their related stimuli (or actions) located on the path towards the preferred state. This leads to Simulation 2 below.

### 5.2. Simulation 2: The effect of needing on reward subsystems (wanting, liking, etc.)

This simulation shows that the presence (versus the absence) of (a path to) a reward decreases the entropy of the states that Agent 1 plans to occupy and increases the associated precision, i.e. the certainty that it should occupy these states. Importantly, our Simulation 2 also shows that the decrease in entropy over which state to occupy, and the increase of associated precision, are magnified when Agent 1 is in a more severe state of need (i.e., when the costs of the non-rewarded states are increased) and there is a path to the preferred state. As an example, if this path to the preferred state is signaled by a pavlovian reward cue processed within the dopaminergic subsystem, then the wanting associated with that cue will be magnified. In other words, the more costly (surprising) these states are, the more the agent is certain that it needs to go to the preferred state. This illustrates how need states amplify the wanting (and/or the liking) of stimuli: by reducing entropy and making the agent more confident about what stimulus, response or course of action to select. Need states have cascading effects also on the stimuli and actions in the path towards goal or reward states. When in a severe need state, relevant stimuli, reward cues and actions have a greater role in reducing entropy and increasing the confidence in the selected course of actions (Parr et al. 2022, Holmes, 2022). These relevant stimuli, reward cues and actions are therefore assigned a greater value and a greater “need-generated” salience which, within the wanting subsystem, would neurophysiologically correspond to increased dopaminergic activity that attribute higher incentive salience, i.e. higher wanting, to reward cues (Berridge, 2004).

### 5.3. Strength and weakness of the model

One advantage of using active inference is its natural incorporation of the pervasiveness of need states through the concept of prior preferences. Conversely, the choice of a reinforcement learning model is motivated by the ability to capture the independent nature of reward prediction and needing. In summary, our simulations show that our model using active inference provides a natural way to model the needing as its own system. Traditional models that discuss need related motivation often assume an automatic link between needing and reward prediction (Keramati & Gutkin, 2014) (also see Berridge, 2023). The proposed model presents a more nuanced view, acknowledging the association and dissociation of needing with other subsystems, such as the wanting subsystem, that process external cues. This broader perspective has the potential to explain a wider range of experimental findings, both at the neural and behavioral levels. Indeed, the increase in precision which, in active inference, is interpreted as neuronal gain (Friston, 2010) could possibly be viewed as the (need induced) salience conferred by need states to relevant stimuli (see Chen et al., 2016). In this sense, if we interpret the increase in precision (in our results) as a need induced neuronal gain toward stimuli/events that are treated within some subsystem, it becomes clear how needing can function independently while interacting with other subsystems like wanting, liking, and interoception. For instance, needing can increase precision of stimuli/events treated within other subsystems, such as the liking or interoceptive ones (Cabanac, 2017; Berridge & Kringelbach, 2015; Becker et al., 2019, Bosulu et al., 2022), even in absence of a pavlovian cue that triggers the wanting subsystem (Wassum et al., 2011 Salamone et al., 2018), and this fits well with some recent meta-analytic results of human fMRI data distinguishing between needing and wanting (see Bosulu et al., 2022). Conversely, a specific need can intensify wanting when a relevant Pavlovian reward cue is present, possibly through this increase in (policy) precision. Notably, this can occur through “active inference,” i.e. without (re)learning (or liking) the consequence of that reward in that specific need state (Berridge, 2007; Zhang et al., 2009; Berridge, 2023). We acknowledge that while simulations illustrate the benefits and predictions of the proposed active inference framework of needing, empirical data testing is still required.

### 5.4. Summary and future developments

The proposed model of needing has potential to be extended to broader psychological phenomena in humans (Pool et al., 2016; Stussi & Pool, 2022; Maslow, 1943; Baumeister & Leary, 1995; Maner et al., 2007). This framework could also be applied to drug dependence/addiction, where a crucial question arises: does drug consumption stem from needing, where the drug state becomes embedded as a prior preference over internal states (Turel & Bechara, 2016; O’Brien et al., 2006), or from wanting, driven by excessive dopamine sensitization to drug-related policies leading to the drug (Berridge & Robinson, 2016; O’Brien et al., 2006)? Depending on the case, different brain regions and behavioral therapy approaches may be targeted.

Overall, this study aimed to provide a conceptual model for needing and its interaction with reward subsystems, based on the active inference framework and the embodied pervasiveness hypothesis. However, further work is needed to fully clarify and empirically test the relationships between the abstract notions introduced here and their underlying biological substrates. A systematic mapping between the information-theoretic concepts used and neurobiological evidence remains an open objective for future research.

## CONFLICT OF INTEREST

The authors declare that they have no conflict of interest.

## ACKNOWLEDGMENTS

The research was supported in part by NSERC Discovery Grant #RGPIN-2018-05698 to SH and UdeM institutional funds to JB and SH. This research is supported by funding from the European Union’s Horizon 2020 Framework Programme for Research and Innovation under the Specific Grant Agreement No. 945539 (Human Brain Project SGA3) to GP and No. 952215 (TAILOR) to GP, and the European Research Council under the Grant Agreement No. 820213 (ThinkAhead) to GP.

## AUTHOR CONTRIBUTION

We thank Kent C. Berridge for providing valuable insights and feedback during the development of this paper.

**Juvenal Bosulu**: Designed the study, performed the experiment, data analysis, interpretation, and wrote the manuscript. **Giovanni Pezzulo**: revised the manuscript and provided critical feedback. **Sébastien Hétu**: revised the manuscript and provided critical feedback. All authors contributed to and approved the final manuscript version.

## DATA AVAILABILITY STATEMENT

All Data is available upon request.

## APPENDIX: SIMULATION AGENTS

### Agent 1

Agent 1 incorporates the main ideas and equations discussed in this article about needing and wanting systems. It is a simplified version of active inference, in which the perceptual part is kept as simple as possible, by assuming that all the states of the grid world are observable. Technically, this means that we are dealing with a Markov Decision Process (MDP) and not a Partially Observable Markov Decision Process (POMDP) as more commonly done in active inference (see Friston et al., 2009; Friston et al., 2017). This simplifying assumption is motivated by the fact that our focus in this work is on action selection and not perceptual discrimination..

In Agent 1, the need state has a pervasive effect on all the states of the grid-world (except the reward state) as described above, in . Following active inference, this pervasive effect is reflected in the prior preferences for the states to occupy (denoted as C, to follow the usual notation of active inference in discrete time), which is greater for the rewarding state than for any other state.

The prior preferences followed a normal distribution centered on the value of the preferred state with a degree of sensitivity regarding deviations from that centered value. The center, i.e. prior preference, is the mean and the sensitivity is the variance, and their values were 1 and 0.5, respectively. (Note that the agent may have behaved differently if the variance/sensitivity was different). This followed the logic of the “homeostatic reinforcement learning” model of Keramati and Gutkin (2014) which is based on minimizing the sum of discounted deviations from a setpoint.

In order to model the pervasive effect of the need state, we assume a joint probability of 1 between the (increasing) need state and other states, except the preferred/satiety state which had a joint probability of 0 with the need state. (Please note that the behavior of the agent may have been different if the joint probability with the other states was significantly less than one.)

Given the pervasiveness of the need state, almost all of the *y*_*i*_ acquire the negative valence and their conditional probability based on prior preference (of satiety) decreases, because their joint probability, i.e. co-occurrence, with satiety decreases. Hence, only the state (or group of states) on the path to the preferred state has high probability as it co-occurs with the prior preference.

Since the agent expects to occupy (or to move towards) these a-priori probable states, the prior over states also translates into priors over actions or action sequences (policies) that achieve such states. In this simplified setting, action (and policy) selection simply corresponds to inferring a distribution of states that it prefers to occupy and policies to reach (sequences of) these states. In other words, Agent 1 tends to select policies that lead it to achieve goal states - which in Bayesian terms corresponds to maximizing model evidence (Parr et al 2022).

#### Agent 2

Agent 2 had the “need state” (state 8) but did not have the pervasive effect of need states as Agent 1. To implement an agent that is guided by reward prediction, we use the reinforcement learning framework. The goal of Agent 2 is to maximize a reward function, through reward prediction. Thus, the agent makes decisions based on prediction of rewards assessed by state-action values, i.e. each decision to pursue a course of actions will depend on the value of actions given the current states (see Sutton & Barto, 2018). Here the policies depending on the action values are denoted *Q*^π^ (*s, a*), and given by:

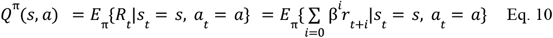

The equation shows the value or “quality” (Q) of the action (a) in state (s) under a policy (π). The function denoted *Q*^π^ (*s, a*) expresses the expected (E) return (R), which is the (expected) discounted (β^*i*^) sum of rewards (*r*_*t*+*i*_), starting from state (s) and taking the action (a), and thereafter following policy (π). Here the state *s* for Agent 2 is equivalent to the state/observation *y* of Agent 1.

Learning of action is updated by the Temporal Difference between the previous action value and the current one as in the equation below:

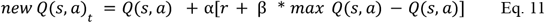

where α represents a learning rate.

The agent’s decision after learning is based on the optimal policy π_*_, i.e. the one that maximizes the expected return, and therefore the optimal *Q*^π^ (*s, a*), noted *Q* ^*^ (*s, a*)is equal to:

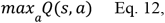

where *max* is related to the action that maximizes *max Q* (*s, a*)

For a better comparison with the agent 1 (that embodied pervasiveness), the actions of the agent 2 were also transformed into a softmax function:

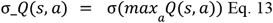

where σ is the softmax function that turns action values into probabilities.

We should keep in mind that agent 2 is used only to illustrate prediction in its simplest form, and does not include advanced reinforcement learning techniques.

## Notes

### Competing Interest Statement

The authors have declared no competing interest.

### Summary of Updates

The manuscript has been modified to focus on the description of the needing mechanism.

